# High-throughput laboratory evolution of *Escherichia coli* under multiple stress environments

**DOI:** 10.1101/143792

**Authors:** Takaaki Horinouchi, Shingo Suzuki, Hazuki Kotani, Kumi Tanabe, Natsue Sakata, Hiroshi Shimizu, Chikara Furusawa

## Abstract

Bacterial cells have a remarkable capacity to adapt and to evolve to environmental changes. Although many mutations contributing to adaptive evolution have been identified, the relationship between the mutations and the phenotypic changes responsible for fitness gain has yet to be fully elucidated. For a better understanding of phenotype-genotype relationship in evolutionary dynamics, we performed high-throughput laboratory evolution of *Escherichia coli* under various stress conditions using an automated culture system. One measure of phenotype, transcriptome analysis, revealed that the expression changes which occurred during the evolution were generally similar among the strains evolved in the same stress environment. We also found several genes and gene functions for which mutations were commonly fixed in the strains resistant to the same stress, and whose effects on resistance were verified experimentally. We demonstrated that the integration of transcriptome and genome data enables us to extract the mechanisms for stress resistance.

**Author summary:** Understanding the relationship between phenotypic and genetic changes is a fundamental goal in evolutionary biology, which can provide insights into the past and future evolutionary trajectories. Evolution of microorganisms in a laboratory has been the primary approach to clarify the mappings of phenotypic and genotypic changes. Here, we performed high-throughput laboratory evolution with bacteria using an automated culture system, to quantify phenotypic and genotypic changes occurred under various stress conditions. We identified various stress-specific gene expression changes and mutations, and contributions of them to fitness gain were validated. These results demonstrated that the integration of phenotypic and genotypic changes makes it possible to extract the mechanisms for stress resistance evolution, which will contribute to bioengineering applications.

## Introduction

Laboratory evolution of microorganisms is a powerful approach for elucidating the nature of evolutionary dynamics[1,2]. Recent advances in measurement technologies including high-throughput sequencing have enabled us to quantify phenotypic and genotypic changes during laboratory evolution, which have provided valuable information on the mechanisms and principles of adaptive evolution[3–5]. The impact of laboratory evolution has extended beyond the field of evolutionary biology into engineering and medicine. For example, by using laboratory evolution approaches, a number of candidate mutations that can contribute to antibiotic resistance have been identified, which sheds light on how to control the emergence of antibiotic resistant strains[6–9]. Laboratory evolution has also become a widely used tool for bioengineering applications[10–13], in order to generate cells with improved growth, production titer, and stress tolerance, which are essential to improved industrial production by microorganisms.

A central problem in laboratory evolution of microorganisms is identification of phenotypic and genotypic changes responsible for an observed fitness gain in an evolved strain. Such identification is often difficult, even after whole-genome sequencing analysis of the evolved strains. In some cases, a large number of genotypic changes, mutations, are fixed in evolved strains, and only a portion of such mutations contribute to the fitness gain. Furthermore, various different mutations can cause a similar phenotypic change[5,9,14], and one mutation can affect various cellular functions. The many-to-many relationships between phenotypic and genotypic changes make identifying the mechanisms of fitness gain challenging.

One strategy to overcome such complexity in the analysis of laboratory evolution is to integrate changes of genomic sequence with various levels of phenotypic data. Such phenotypic data include those from omics analysis such as transcriptome and metabolome. Based on such phenotypic data, one can extract correlated phenotypic changes to the observed fitness gain, which facilitates identification of mutations responsible for the phenotypic and fitness changes. For example, analysis of laboratory evolution of antibiotic resistant *E. coli* revealed that the change in antibiotic resistance can be quantitatively predicted by the transcriptomic change during adaptive evolution, which led to identification of genomic mutations contributing the resistance[9]. Furthermore, cross-stress protection, a phenomenon whereby stress resistance is acquired to one stress after adaptive evolution to another stress, also provided valuable information on which phenotypic and genotypic changes related to the stress resistances when analyzed by integrating transcriptome and genome resequencing data[15]. Integration of various levels of phenotypic data with the change of genomic sequence is the future of investigating evolutionary dynamics and will be an essential method for the application of laboratory evolution in medical and engineering fields.

Given the importance of the integration of multiple levels of phenotypic and genotypic data in laboratory evolution, we performed high-throughput laboratory evolution of *E. coli* cells under 11 different stress conditions and then quantified phenotypic changes, specifically transcriptome expression changes and growth rate, and genotypic mutations of independently evolved strains. Transcriptome analysis revealed that, the expression changes which occurred during the adaptive evolution were generally similar among the strains evolved in the same stress environment. We analyzed the correlation between these expression changes and mutations commonly fixed in the strains resistant to the same stress. This analysis revealed the mechanisms for some of the stress resistances, which were validated by introducing the relevant mutations into the genome of the parent strain. The integration of the multilevel phenotypic and genotypic data enables us to understand a wide spectrum of molecular mechanisms in *E. coli* stress resistances.

## Results

### Laboratory evolution under 11 stress conditions and analysis of cross-protection

We selected 11 stresses that cover a wide range of environmental stresses (Table 1). *E. coli* MDS42 cells[16] were cultured in 200 μL of M9 synthetic medium[17] with constant concentration of stressors. The concentrations of these stressors were set to levels which initially decreased the specific growth rate by approximately one half of the non-stress growth condition. Every 6 hours, a fraction of the cells was transferred into fresh medium with the stressor. The transfer volume was adjusted to keep the final cell concentration below a threshold and to maintain the culture in exponential growth phase. The specific growth rate was quantified by initial and final cell concentration, which was used as the measure of fitness under these stress environments (Fig 1a). To evaluate the reproducibility of the evolutionary pathways, for each stress, five independent culture lines were propagated in parallel. For this high-throughput laboratory evolution, we used a custom-developed automated system for laboratory evolution[18], by which we can maintain hundreds of independent culture series in a fully automated manner. After 906 hours of propagation, we observed significant increases in specific growth rate in all 11 stress environments (Fig 1b to 1m). In addition to the cultures with stressors, we maintained the cells in the synthetic medium without adding stressors (Fig 1n). In these cultures, without stress, slight increases of specific growth rate were also observed, although these increases were significantly smaller than those in the stress conditions. For all resistant strains, we confirmed maintenance of the stress resistances by reintroduction after cultivation in the absence of the stressors for at least 60 generations, indicating that the phenotypes of stress resistance were stable.

**Figure 1.**
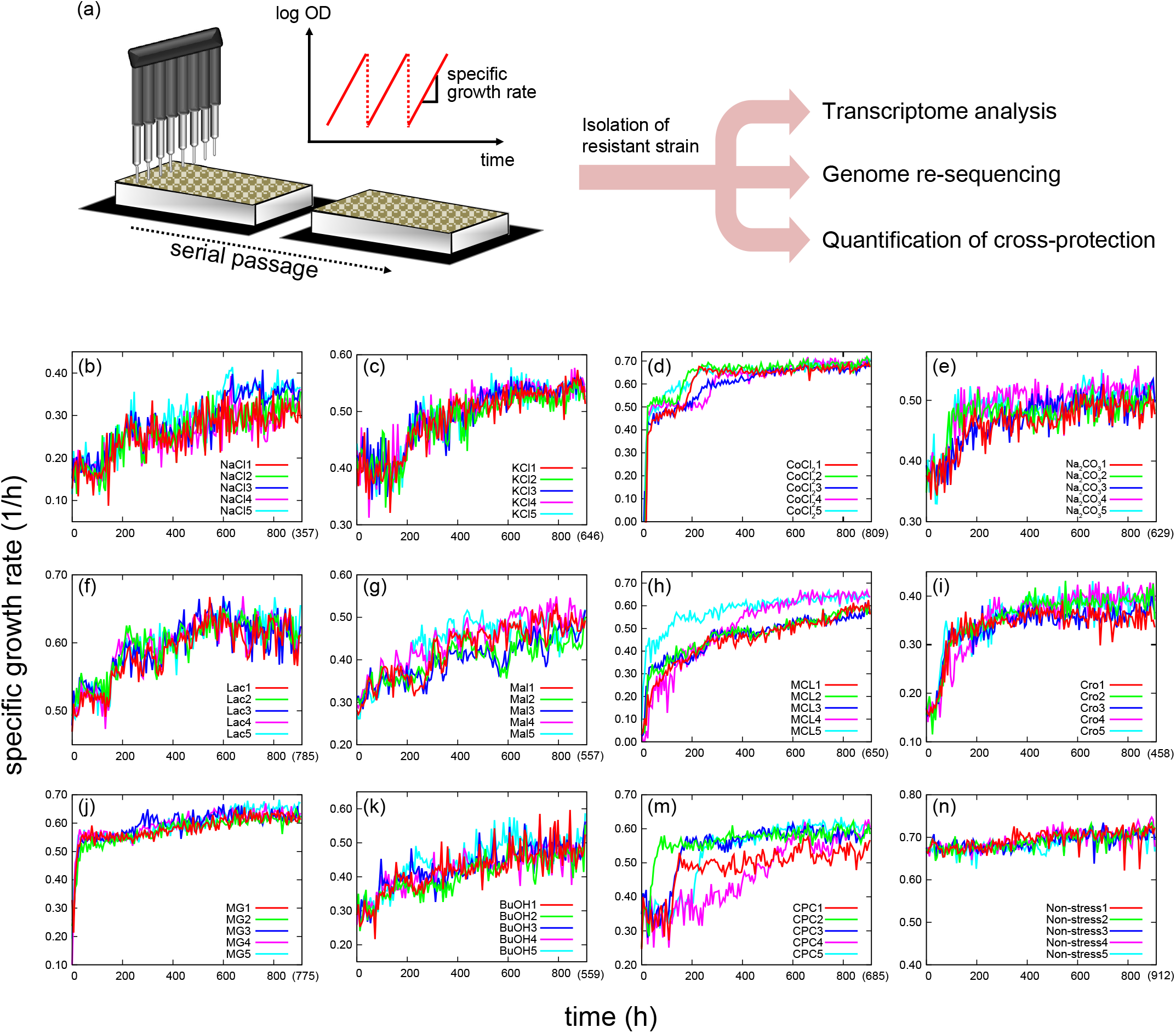
Laboratory evolution under environmental stresses. (a) Overview of the experimental setting. *E. coli* cells were evolved in 11 stress environments by using the automated culture system. Selected clones from adapted populations were sequenced and transcriptional profiles were quantified to analyze phenotype-genotype relationships. (b)-(n) The time courses of the specific growth rate in experimental evolution. Five parallel series of experiments were performed. The values in parentheses show the average number of generations at the end-point of cultivation.

**Table I.** List of stress environments used for laboratory evolution

| **Stressor** | **Abbreviation** | **Concentration** | **Description of stress** |
| --- | --- | --- | --- |
| Sodium chloride | NaCl | 400 mM | Inorganic salt |
| Potassium chloride | KCl | 210 mM | Inorganic salt |
| Cobalt chloride | CoCl<sub>2</sub> | 16 µM | Heavy metal |
| Sodium carbonate | Na<sub>2</sub>CO<sub>3</sub> | 32.5 mM | Alkali |
| L-Lactate | Lac | 40 mM | Organic acid |
| L-Malate | Mal | 30 mM | Organic acid |
| Methacrylate | MCL | 8.75 mM | Unsaturated carboxylic acid |
| Crotonate | Cro | 50 mM | Unsaturated carboxylic acid |
| Methylglyoxal | MG | 350 µM | Ketoaldehyde |
| *n*-butanol | BuOH | 1.25 % | Alcohol |
| Cetylpyridinium chloride | CPC | 4.8 µM | Surfactant |

To explore evolutionary cross-protection, for each obtained resistant strain, we measured the specific growth rate under all 11 stress conditions. In Fig 2, the relative growth rates of the resistant strains to the parent strain are presented (All the growth rates are presented in S1 Table). The results demonstrated that the stress resistant strains often show cross-protection and hyper-sensitivity to multiple stresses. A clear symmetric cross-protection was observed among NaCl and KCl stress resistant strains, i.e., NaCl resistant strains exhibited significant KCl resistance, and vice versa. This fact suggested that, at least a part of the mechanisms of the resistances were shared among NaCl and KCl resistant strains. For other cases, the cross-protections were asymmetric, which suggested that the mechanisms of these resistance and hyper-sensitivity were independent, or there are hierarchical relationships between the resistance/hyper-sensitivity mechanisms. For example, methylglyoxal (MG) resistant strains exhibited resistance to lactate (Lac), while Lac resistant strains did not significantly affect MG resistance. This asymmetric cross-protection might be explained by a hierarchical structure of the resistance mechanisms. To degrade MG in metabolic pathways, one possible intermediate is Lac[19], and thus MG resistance strains might need to have Lac resistance at the same time. In contrast, MG is not involved in Lac the degradation pathway, and thus Lac resistance would not necessarily be accompanied with MG resistance.

**Figure 2.**
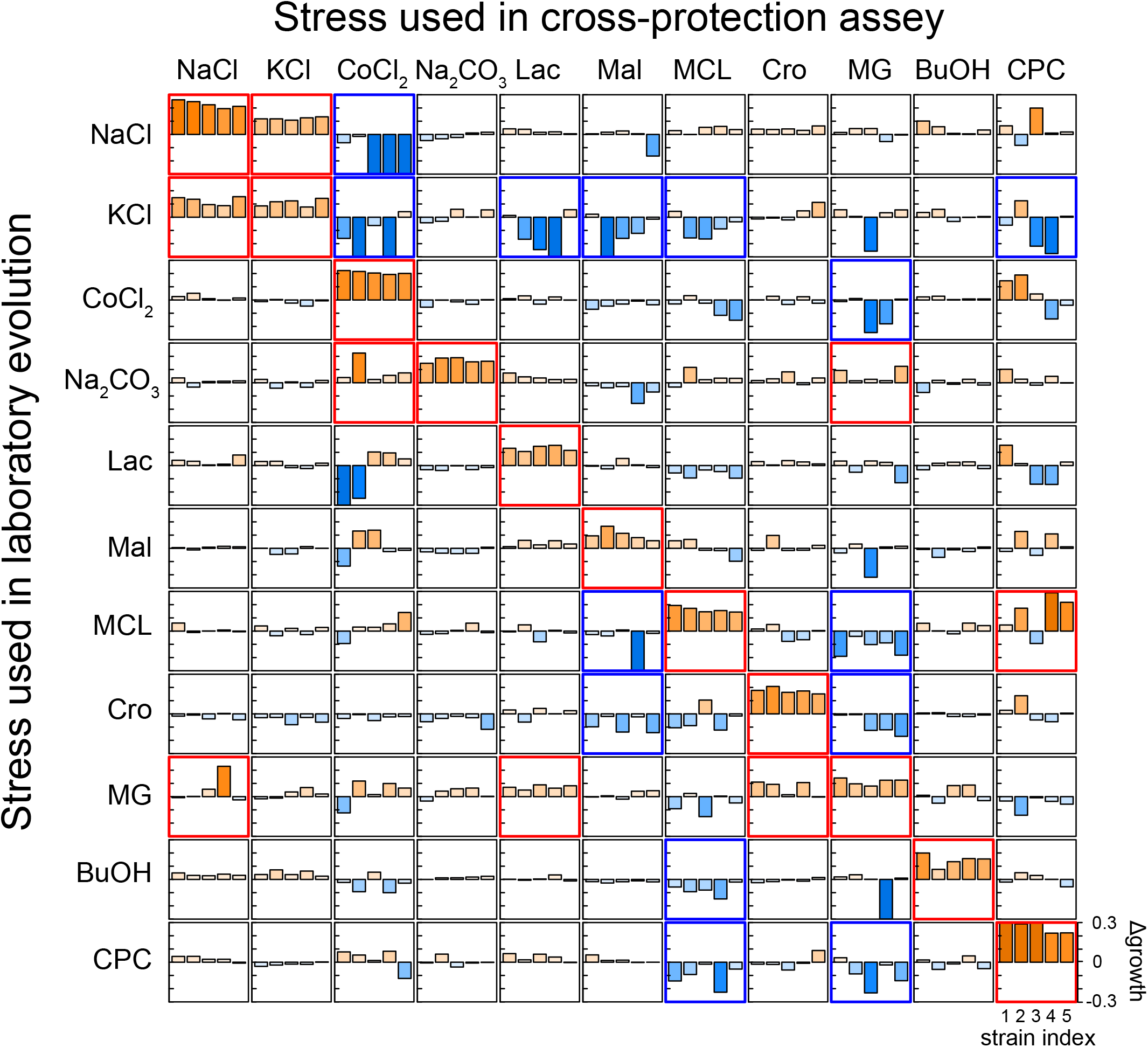
Quantification of cross-protection and hyper-sensitivity. The vertical axis of each figure shows the difference of specific growth rate compared to the parent strains, while the horizontal axis corresponds to the index of five resistant strains obtained independently. Each column of figures corresponds to the data obtained under a same stressor, while each row indicates the stress used for the laboratory evolution.

### Changes in expression profiles in stress resistant strains

To elucidate the mechanisms of the various stress resistance phenotypes, we performed transcriptome analysis of all the resistant strains we created by using microarray experiments (all transcriptome data are presented in S2 Table). The results demonstrated that the expression changes which occurred during the long-term cultivation were generally most similar among the strains evolved in the same stress environment, as shown in the hierarchical clustering analysis transcriptome data (Fig 3a). To characterize the similarity in transcriptomic changes in these resistant strains, the expression changes of representative transcriptional factors (TFs) are presented in Fig 3b. As shown, these TFs exhibited similar expression patterns within each stress resistant clade. For example, expression levels of *marR* encoding a repressor protein for the *marRAB* operon were commonly down-regulated in BuOH and Na_2_CO_3_ resistant strains (S1a Fig). The *marRAB* operon is involved in resistance to several stresses including antibiotics and organic solvents[20], and the inactivation of *marR* has been shown to increase the resistance of *E. coli* to multiple antibiotics and organic solvents by up-regulation of the AcrAB-TolC multidrug efflux pump[21,22]. Our results also showed a negative correlation between *marR* and *acrAB* expression levels (S1b Fig), which suggests up-regulation of *acrAB* contributed to the observed BuOH and Na2CO3 resistances. In contrast, *marR* expression was up-regulated in Crotonate (Cro) resistant strains. Although this characteristic up-regulation might be involved the resistance, the mechanism remains unclear. For another example, *hns* encoding a histone-like protein H-NS were up-regulated in all CoC, Lac, Mal, MG, and CPC resistant strains (S2a Fig). H-NS is a global regulator involved in the adaptation to various stress environments, whose change in expression level affects hundreds of genes. Among these target genes, we found that *rcsD* expression levels in the resistant strains clearly showed a negative correlation to *hns* expression level (S2b Fig). *rcsD* is known to be involved in acid stress resistance[23], and thus the correlation suggests the contribution of *hns* regulation to stress resistance through *rcsD* expression changes. The observed similarity of expression changes in the resistant strains, as represented by the TFs expression patterns in Fig 3b, strongly suggests that such expression changes contributed to the acquisition of the resistances.

**Figure 3.**
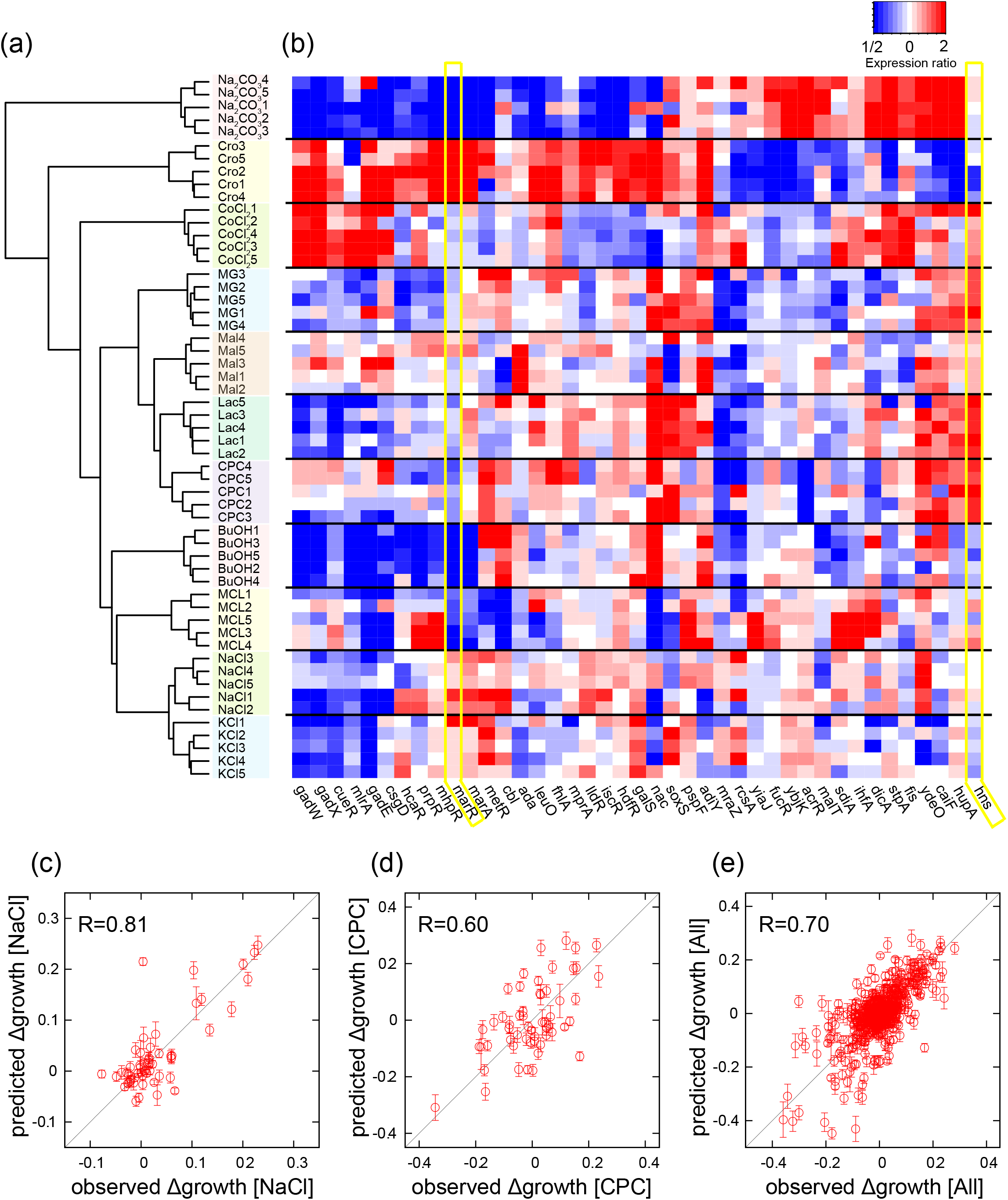
Transcriptomic changes in resistant strains. (a) Hierarchical clustering of overall expression changes in the resistant strains. The expression changes were calculated by dividing expression level of genes in the resistant strains with that of the parent strains, where these expressions were quantified under the condition with the corresponding stress addition. (b) Expression changes of transcriptional factors. Representative transcriptional factors having larger variance of expression changes over resistant strains were plotted. The expression changes more than 2-fold or less than 1/2 are shown in the same color with 2 and 1/2, respectively. (c-e) Prediction of growth rate using transcriptomic changes. Comparisons between observed and predicted growth rate under (c) NaCl, (d) CPC, and (e) all data were calculated by fitting using the following 15 genes: *tbpA, appB, ydiH, gadE, sbp, aldB, asr, marC, proW, tktB, nac, thiC, ydhZ, acs*, and *gcd*. Only test data, not used for the fitting, are plotted. The error bars in the y-axis were obtained by predicted growth rate calculated from 10,000 different sets of test data and training data. For the details of the prediction of growth rate, see Materials and Methods.

To further analyze the correlations between two elements of phenotype, transcriptome changes and stress resistance, we used a simple, previously published[9] mathematical model to predict the resistances using the obtained gene expression profiles (see Materials and Methods for details). Briefly, we assumed the changes in the specific growth rates under stresses are represented by a linear combination of log-transformed expression changes during the long-term cultivation. Then, we sought the optimal number and combination of genes which have the highest predictive accuracy by using cross-validation and genetic algorithm. As result, we found that the combinations of 15 to 20 genes offered the highest prediction accuracy on average (S3a Fig). Fig 3c-e show examples of prediction accuracy by the linear model with 15 genes (see legend of Fig 3 for details; all fitting results are shown in S3 Fig). In this analysis, for each stress environment, the coefficients of the linear model were estimated by fitting the training data, while the plotted data are test data that were not used for fitting. The estimated growth rate under stresses agreed with those observed, indicating that this linear model can predict the change of stress resistance phenotype by a relatively small number of genes. This analysis enabled us to extract the genes whose expression changes provided the most relevant information for predicting stress resistance (S3k Fig). For example, *tbpA* encoding thiamin ABC transporter was specifically up-regulated in Na_2_CO_3_ resistant strains, while it was down-regulated in NaCl, KCl, and Cro resistant strains (S4a Fig). Similarly, *ydiH* which encodes a predicted protein with unknown function was commonly up-regulated in CoCl_2_ and Cro resistant strains, while it was down-regulated in several resistant strains (S4b Fig). Here, we successfully screened genes whose expression changes are highly correlated to resistance acquisition, which can contribute to highly accurate descriptions of complex evolutionary dynamics.

### Mutations fixed in stress resistant strains

The mutations identified in the resistant strains are shown in Fig 4, and the detailed information about the mutations is presented in S3 Table. Less than 10 mutations were fixed in each of the resistant strains. We also sequenced two strains maintained without addition of stress (Fig 1n), and found that relatively small number of mutations were fixed in these control strains, as shown in Table S3.

**Figure 4.**
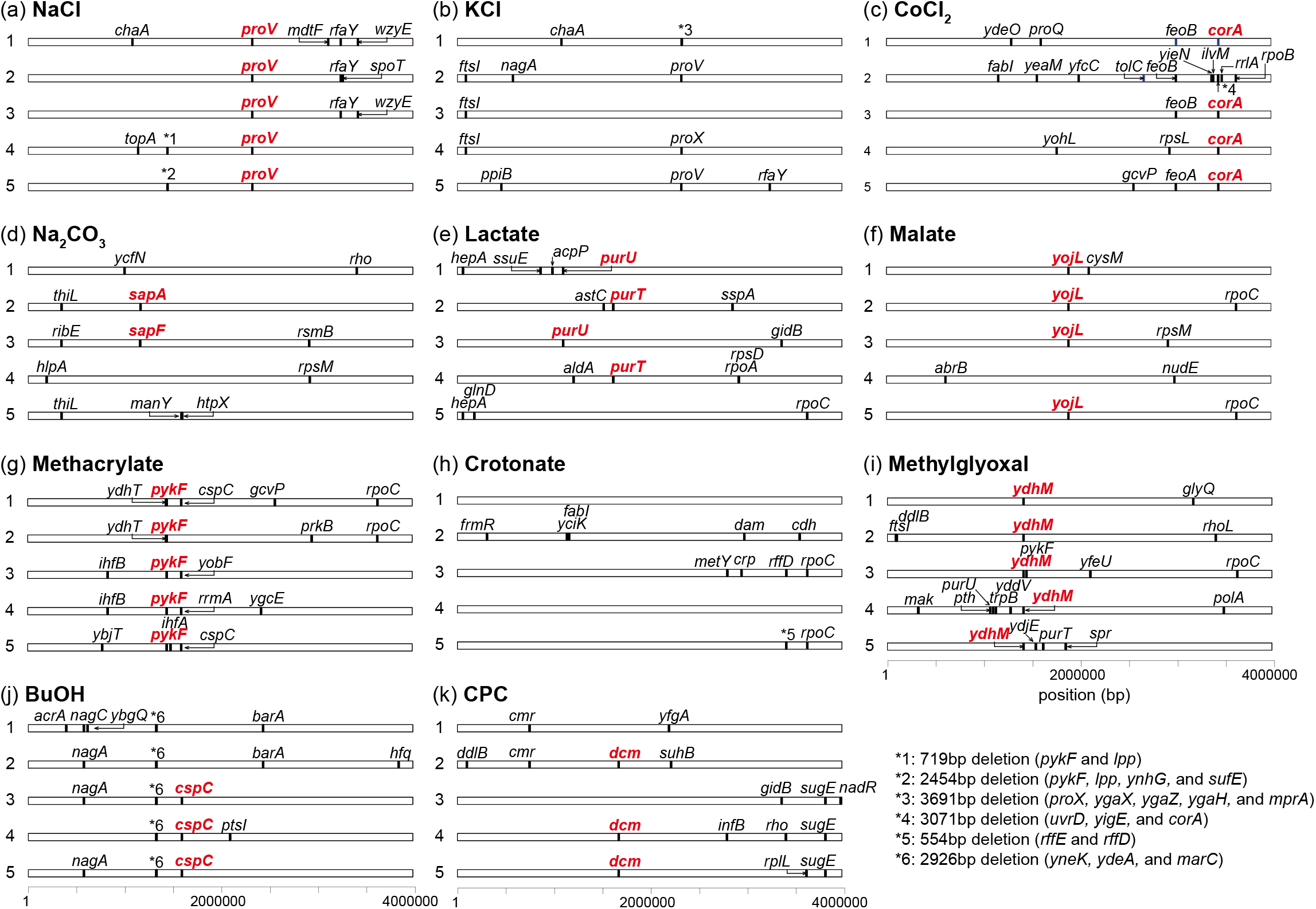
The map of mutations identified in the resistant strains. Coordinates are relative to the reference MDS42 genome. The representative mutations commonly identified in the resistant strains to a same stress are highlighted by red characters, and whose effects on the corresponding stress resistance were evaluated as shown in Fig 5. The detailed information on the identified mutations was shown in S3 Table.

We found several genes and gene functions for which mutations were commonly fixed in the resistant strains, suggesting contributions by these mutations to the resistances. To verify the possible contribution of these mutations to the stress resistance, we introduced some of such commonly identified mutations (highlighted by a red character in Fig 4) into the genome of the parent strain, and quantified the change of growth rate under corresponding stresses (Fig 5). Below, we discuss some examples of the relationship between resistance acquisition and phenotype/genotype changes. A full description of the discussions is presented in S1 Text.

**Figure 5.**
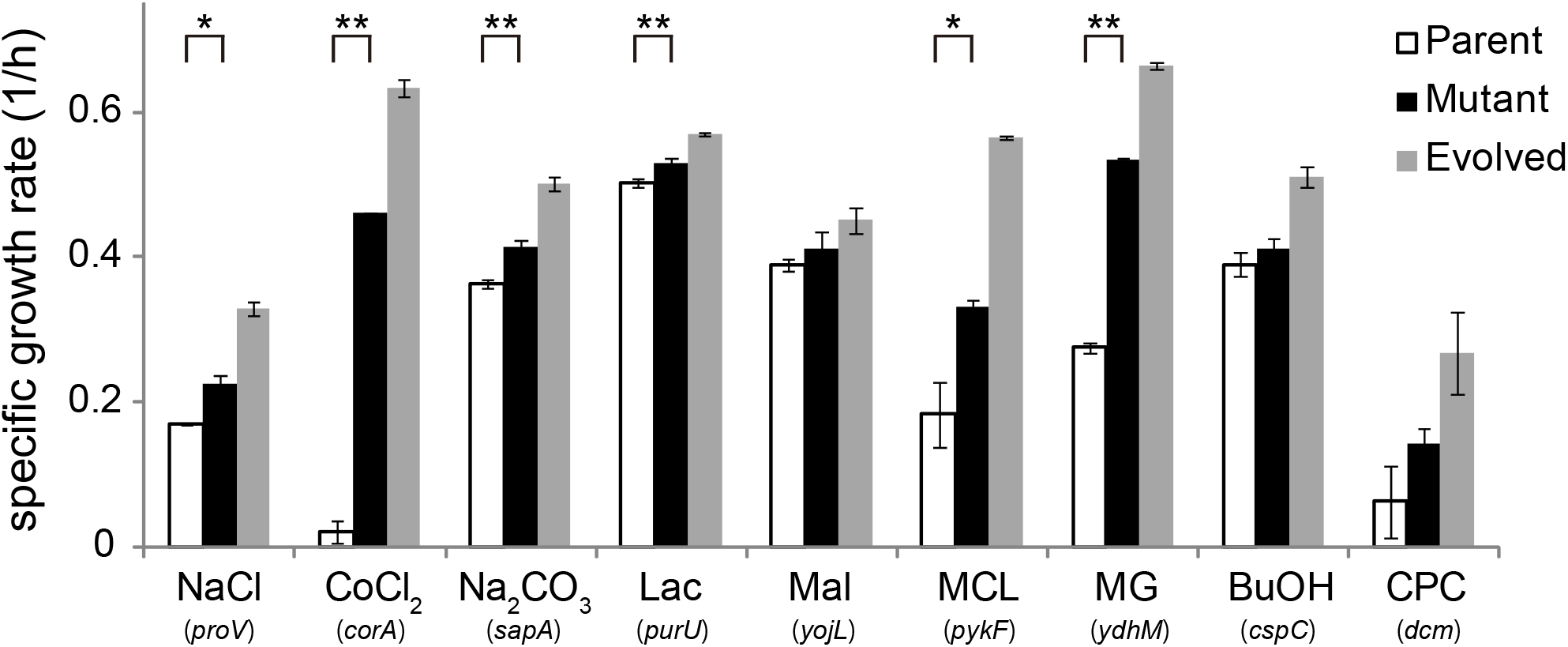
Growth rates of site-directed mutants. One of mutations commonly identified in the resistant strains to a given stress (highlighted by red character in Figure 4) were introduced back to the parent strain. For each mutant, the names of the mutated genes are shown. Error bars indicate standard deviations calculated from three independent cultures. Statistical analysis was performed by t test. Significances were accepted at \**P* < 0.05, and **P <0.01.

The first example of the common mutations is that of to the *proU* ABC transporter which was identified in all NaCl resistant strains and 4 KCl resistant strains (Fig 4a and 4b). All of these mutations caused frame-shifts, suggesting that they disrupted the activity of the *proU* transport system. The *proU* operon (*proVWX*) encodes a binding protein-dependent transport system which is essential for the uptake of osmoprotectants such as glycine betaine and is known to be up-regulated in response to osmotic stress[24]. Our expression analysis showed that, the expression of *proU* operon was significantly up-regulated (> 50 times) in response to the initial NaCl stress addition, however after long-term cultivation the expression levels decreased when the cells acquired NaCl resistance (S5 Fig). These results suggested that, the programmed up-regulation of *proU* operon in response to osmotic stress is not beneficial in an environment without osmoprotectants such as that which we used. Instead, the up-regulation of *proU* operon suppresses the cellular growth presumably due to its energy consumption. Thus, the disruption of *proU* transport activity can contribute to the active cellular growth under NaCl stress. To support this hypothesis, we introduced a frame-shift mutation identified in one of the NaCl resistance strains into the parent strain using site-directed mutagenesis, and confirmed that this mutation increased the growth under NaCl stress (Fig 5).

For another example, all resistant strains to MG stress had mutations related to *ydhM* (*nemR*) as shown in Fig 4i, which encodes N-ethylmaleimide reductase repressor (4 strains had mutations in the upstream region and one strain had mutation in the coding region of *ydhM*). We confirmed that the introduction of mutation in the upstream region of *ydhM* caused significant increase of growth rate under MG stress (Fig 5). The expression level of *ydhM* was significantly up-regulated (approximately 100 times) by the addition of MG stress to the parent strain, while similar expression levels were maintained after the acquisition of MG resistance by laboratory evolution (S6a Fig). Interestingly, after the adaptive evolution to MG stress, the up-regulation of *ydhM* was observed even without addition of MG stress (S6a Fig), which suggested that the identified *ydhM* mutations in the resistant strains serve to “assimilate” the stress response into a genetically encoded invariant resistance phenotype. This genetic assimilation[25,26] might contribute to the robustness of the resistance phenotype by altering reaction norm[27] to MG stress. *ydhM* expression controls the expression of *gloA*, which is involved in the degradation pathway of MG to lactate and is known to contribute to MG resistance[19]. There is a clear correlation between *ydhM* and *gloA* expression levels in the resistant strains (S6b Fig), which suggests that the observed MG resistance was caused by increasing MG degradation activity through the up-regulation of *gloA*.

The detailed descriptions for the other common mutations found in the resistant strains are presented in S1 Text. We demonstrated that the identified common mutations generally contributed to resistance acquisition, as shown in Fig 5. It should be noted that, to understand the mechanism how such mutations affected the stress resistance phenotype in addition to the identification of genomic mutations, the information on expression changes, transcriptome phenotype, during the adaptive evolution was also highly valuable, as in the cases of *proU* and *ydhM* mentioned above.

It is worth noting that, although in some cases the data suggested simple causal relationships between genomic mutations and phenotypic changes, the relationships between them were not always a simple one-to-one correspondence. As shown in Fig 3, the expression changes of various genes were generally similar among independently evolved resistant strains to a given stress, while the number of fixed mutations was often small and the overlap among them was imperfect. For example, the expression changes of TFs in five Na_2_CO_3_ resistant strains are quite similar during their adaptive evolution as shown in Fig 3b. In contrast, there was no gene or gene function for which mutations were fixed in common in all the Na_2_CO_3_ resistant strains. One possible explanation for these results was that mutations in different genes caused similar expression changes. Since it is known that epistatic interactions among mutations are ubiquitous, similar expression changes caused by different mutations might emerge through selection under stresses. Another possibility is that epigenetic factors without alternation of genome sequence might contribute to the stress resistance acquisition, as suggested in several previous studies[28,29].

## Conclusion

Laboratory evolution experiments combined with organism-wide phenotype and genotype analyses enable us to understand mechanisms of adaptive evolution. In this study, we demonstrated that the high-throughput system for laboratory evolution we developed can create resistant strains of *E. coli* to various stress environments by long-term cultivation. Using this system, we can cultivate up to 44 microplates (96 or 384 wells) simultaneously, and thus more than 10,000 independent culture series can be maintained in a fully automated manner. This system allows us to trace evolutionary dynamics under various environmental conditions and initial conditions (e.g., all *E. coli* strains in single-gene knockout library[30]), with numerous replicate experiments.

In this system for laboratory evolution, we maintain cells under relatively small culture scale (i.e., 200 μL). One might question whether such small culture scale might result in small population size, which causes fixation of random drifts and accumulation of neutral mutations. Our data suggested that this is not the case. We confirmed that the population size was maintained more than 10^5^ cells. The ratio of synonymous to non-synonymous substitutions over all resistant strains was relatively small in comparison with that of neutral mutations, suggesting that the fixation of a majority of substitutions were driven by evolutionary selection pressure. The fact that, for many stresses, resistant strains to a common stress shared common mutations also supports the idea of evolutionary selection.

The genome-wide expression and resequencing analyses showed common phenotypic and genotypic changes in the resistant strains to an identical stress. It is worth noting that, our results demonstrated that combined use of transcriptome and genome resequencing analysis greatly accelerated our interpretation about how *E. coli* strains acquired the stress resistance. For example, mutations related to the *proU* operon found in all NaCl resistant strains were thought to cancel the deleterious regulatory program. Using only sets of fixed mutations in resistant strains often makes it difficult to establish the causal relationship with stress resistance. Integrating both gene expression and genotype data can overcome this problem and it allows a highly accurate description of the mechanisms responsible for stress resistance.

The adaptive evolution to environmental changes is a phenomenon that involves changes in genome, transcriptome, metabolome, and so on, meaning a complex interaction network is involved. One possible strategy to understanding such complex dynamics is to analyze large-scale data for each hierarchical layer and then to integrate the analyses to extract the essential components for the phenotypic and genotypic changes. The present study is one of the first to achieve genome-wide analysis of evolved strains obtained by high-throughput laboratory evolution, and we succeeded in extracting phenotypic/genotypic changes responsible for stress resistance. Such large-scale data of laboratory evolution will, we believe, help us to unveil principles of evolutionary dynamics and provide valuable information on rational design of industrially useful microbial strains.

## Materials and Methods

### Laboratory evolution

The IS elements-free *Escherichia coli* strain MDS42[16] was purchased from Scarab Genomics and used as the parent strain of the laboratory evolution. The use of the IS elements-free strain can make the resequencing analysis reliable since the determination of the precise position of IS element insertions is often difficult by using high-throughput sequencers. For serial transfer culture, we used 200 μL modified M9 medium[17] including 5g/L glucose as a carbon source. The cells were cultured in 96-well microplates (Corning Inc. 3595) with shaking at 300 rotations/min at 34 °C. All cultivations were performed by the automated culture system[18] (Fig 1a) which consists of a Biomek^®^ NX span8 laboratory automation workstation (Beckman Coulter, Tokyo, JP) in a clean booth connected to a microplate reader (FilterMax F5; Molecular devices, Tokyo, JP), shaker incubator (STX44; Liconic, Mauren, LI), and microplate hotel (LPX220, Liconic, Mauren, LI). The movie of this automated culture system for laboratory evolution is available at youtube (https://www.youtube.com/watch?v=4k6qCN7ppsk). We diluted the cells into a fresh medium every 6 hours. The cells were maintained in the exponential growth phase by adjusting the initial cell concentration of each dilution to keep a final cell concentration of less than 10^7^ cells per well as measured by optical density at 620 nm (OD_620_). Before laboratory evolution under the stress conditions, we cultivated the cells without adding stressors for 96 hours (approximately 90 generations) to adapt them to M9 medium. The specific growth rate was calculated based on the initial and final cell concentrations of the each dilution. The cells after the evolution experiments were single-cloned and were stored as glycerol stocks at −80 °C and used for further analysis. The quantifications of specific growth rates of evolved or genetically manipulated strains (Fig 2 and Fig 5) were performed after 60 hours preculture (approximately 30 to 60 generations) while maintaining exponential growth phase. For the preculture, the same culture conditions as the laboratory evolution with the corresponding stressor were used.

### Transcriptome analysis by microarray

The cells were precultured for 60 hours under the same condition with the laboratory with or without the corresponding stressors. Then, 5×10^7^ cells in the exponential growth phase were killed immediately by the addition of an equal volume of ice-cold ethanol containing 10% (w/v) phenol. After that, the cells were harvested by centrifugation and stored at −80 °C before RNA extraction. Total RNA was isolated and purified from cells using an RNeasy mini kit with on-column DNA digestion (Qiagen, Hilden, Germany) in accordance with the manufacturer’s instructions. The quality of the purified RNA was evaluated using Agilent 2100 Bioanalyzer with an RNA 6000 Nano kit (Agilent Technologies). The purified RNAs were stored at −80 °C prior to transcriptome analysis. Microarray experiments were performed using custom designed the Agilent 8×60K array for *E. coli* W3110, in which 12 probes were prepared for each gene. One hundred ng of purified total RNAs was labeled using the Low Input Quick Amp WT Labeling Kit (Agilent Technologies) with Cyanine3 (Cy3) in accordance with the manufacturer’s instructions. After confirmation of yields (> 825 ng) and specific activities (> 15 pmol/μg) of the Cy3-labbeled cRNAs using NanoDrop ND-2000, labeled cRNAs (600 ng) were fragmented and then hybridized on the microarray for 17 h while rotating at a speed of 10 rpm at 65 °C in an hybridization oven (Agilent Technologies). Washing and scanning of microarrays were performed in accordance with the manufacturer’s instructions. Microarray image analysis was performed using Feature extraction version 10.7.3.1 (Agilent Technologies). The background corrected intensity values were normalized using the quantile normalization method. The normalized data of microarrays have been deposited in GEO under the accession code [GSE89746] and are presented in S2 Table. In that table, we also present the biological triplicate data to check the reproducibility of the analysis, i.e., expression data obtained from different cultures of the parent strain without addition of stress. So as to use only quantitatively reliable data, genes with low expression levels (less than 100 a.u. in any strain) were excluded from the following analysis (approximately 60 *%* of genes remain). We confirmed that after the removal of low expression genes, more than 99 % of expression ratios between the biological repetitive data were within the range 1/1.3 to 1.3.

### Predicting stress resistance by gene expression levels

To examine the contribution of the gene expression changes to the stress resistances, we constructed a simple mathematical model to predict the resistances using the obtained gene expression profiles[9]. Here, we assumed that the stress resistances, quantified by the change of growth rate after stress addition, are determined as a function of gene expression levels and neglected any direct effect of the mutations on the resistance. Furthermore, for simplification, we neglected non-linear effects and cross terms of the gene expression changes. Thus, we assumed the following simple linear model to predict the growth rate change by the expression levels of *N* genes:

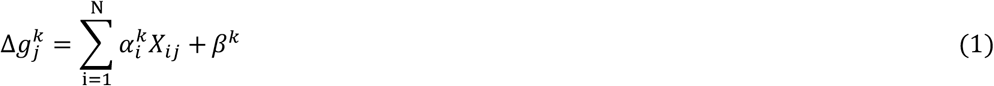

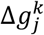 is the growth rate changes in the *j* th strain for the *k* th stress, *X_ij_* is the log_10_-transformed expression level of the *i* th gene in the *j* th strain after standardization to zero mean and unit variance, and 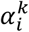 and *β^k^* are fitting parameters. Since the number of genes used in this analysis is much larger than the number of fitness data, the use of all transcriptome data for the fitting resulted in overfitting leading a meaningless prediction of the fitness. In order to avoid such overfitting and to investigate the number of genes with the highest predictive accuracy, we used the cross-validation method, in which the fitness data were separated into training data used for the parameter fitting and test data used to verify the prediction accuracy. When *N* was large, the prediction accuracy for the test data became small due to overfitting, while the accuracy became small when *N* was small since the linear combination of expressions was insufficient to represent changes of the fitness data. In this analysis, we used a 4-fold cross validation method. That is, the resistant strains were randomly partitioned into 4 equally sized subgroups; 1 subgroup was used as the test data set for validation and the remaining 3 subgroups were used for the fitting.

Expression levels of genes in the same operon are generally correlated, which can cause problems in the gene selection procedure. Thus, in each operon, we selected a gene having the highest average expression level over all samples and use it for the fitting, while we discarded the data of other genes. Furthermore, we excluded data with relatively small expression changes among the resistant and parent strains, since the expression changes of such relatively unchanged genes dominated the experimental errors. These selection criteria left 413 genes for the analysis.

*N* genes used for the fitting were selected by a genetic algorithm (GA), in which the correlation coefficient between the predicted and observed growth rate changes of the training data sets was used as the fitness function. As an initial population, 1000 sets of *N* genes were randomly selected, and the fitness of the sets was calculated. Then, gene sets within the top 5% of the highest fitness were selected as the parent sets of the next generation from which mutant sets were generated by randomly replacing a single gene. We iterated 300 cycles of the mutant sets and selected gene sets with the highest fitness to obtain sets of small number of genes whose expression levels can represent changes in drug resistances and susceptibility. We repeated the selection of gene sets using 10,000 different training data sets prepared by randomly partitioning the total data set to obtain the frequency of genes selected after the GA, as shown in S3b Fig.

### Genomic DNA preparation

We prepared precultures by shaking stocked strains in 200 μL of modified M9 medium in 96-well microplates for 23 h at 34 °C without stressors. The precultured cells were diluted to an OD_600 nm_ of 3 × 10^−5^ in 10 mL of fresh modified M9 medium in test tubes. Cell culture was performed at 34 °C for 23 h with shaking at 150 strokes min^−1^ using water bath shakers (Personal-11, Taitec Co.), and we confirmed that the OD_600 nm_ values reached more than 1.0. Rifampicin (final concentration 300 μg/mL) was subsequently added, and the culture was continued for a further 3 h to block the initiation of DNA replication. The cells were collected by centrifugation at 25 °C and 20,000 × *g* for 5 min, and the pelleted cells were stored at −80 °C prior to genomic DNA purification. Genomic DNA was isolated and purified using a Wizard^®^ Genomic DNA Purification Kit (Promega) in accordance with the manufacturer’s instructions. To improve the purity of genomic DNA, additional phenol extractions were performed before and after the RNase treatment step. The quantity and purity of the genomic DNA were determined by the absorbance at 260 nm and the ratio of the absorbance at 260 and 280 nm (A_260/280_) using NanoDrop ND-2000, respectively. As a result, we confirmed that A_260/280_ values of all samples were greater than 1.7. The purified genomic DNAs were stored at −30 °C prior to use.

### Genome sequence analyses using Illumina HiSeq System

Genome sequence analyses were performed with Illumina HiSeq System. An 150 bp paired-end library was generated according to the Illumina protocol and sequenced in Illumina HiSeq. In this study, 58 samples with different barcodes were mixed and then sequenced in two lanes, resulting in about 300-fold coverage on average. The raw sequence data are available in the DDBJ Sequence Read Archive under accession number [DRA005229].

The quality of the sequence data was first assessed with FastX Toolkit 0.0.13.2 (http://hannonlab.cshl.edu/fastx_toolkit), and the raw reads were trimmed using PRINSEQ[31], in which both ends with quality scores lower than Q20 were trimmed. The potential nucleotide differences were also validated with BRESEQ version 0.28[3]. For structural variations, we used the same method as above[5] to detect large ins/del.

### Construction of deletion and single nucleotide substitution strains

To construct mutated strains shown in Fig 5, we introduced identified mutations into the parent strain by the markerless gene replacement method[32]. Briefly, to construct DNA fragments that had deleted coding regions, upper flanking regions of the start codon were amplified by PCR using genomic DNA of the parent strain as templates with forward primers containing the *Eco*RI site and reverse primers containing overlaps with lower flanking regions of the stop codon. The lower flanking regions were amplified by PCR with forward primers containing overlaps with the upper flanking regions and reverse primers containing the *Kpn*I site. After purification by the MinElute PCR Purification Kit (Qiagen), the PCR products were combined by overlap extension PCR. To construct DNA fragments that introduce an identified mutation, DNA fragments were amplified by PCR using genomic DNA of each resistant strain in which the mutation was identified. Each DNA fragment was purified by the MinElute PCR Purification Kit and then cloned into the suicide plasmid pST76-K[32] (The plasmid was kindly provided by Dr. György Pósfai, Biological Research Centre of the Hungarian Academy of Sciences, Hungary). After confirmation of DNA fragment sequences by Sanger method, transformation, integration into the parent strain genome, replacement stimulated by double-stand break, and plasmid curing were performed in accordance with a previous method[32]. After construction of mutant strains, corresponding genomic regions were amplified by PCR and then confirmed by Sanger sequencing of the PCR products directly.

## Acknowledgements

We thank Dr. Kylius Wilkins for proofreading of the manuscript and constructive comments, and Mr. Tatsuya Itoga, Dr. Satomi Banno, and Dr. Katsunori Yoshikawa for technical assistance. This work was supported in part by Grant-in-Aid for Scientific Research (B) [26290071, 15KT0085], Grant-in-Aid for Scientific Research (A) [24246134], and a Grant-in-Aid for Scientific Research (S) [15H05746] from JSPS.

---

**S1 Figure.**
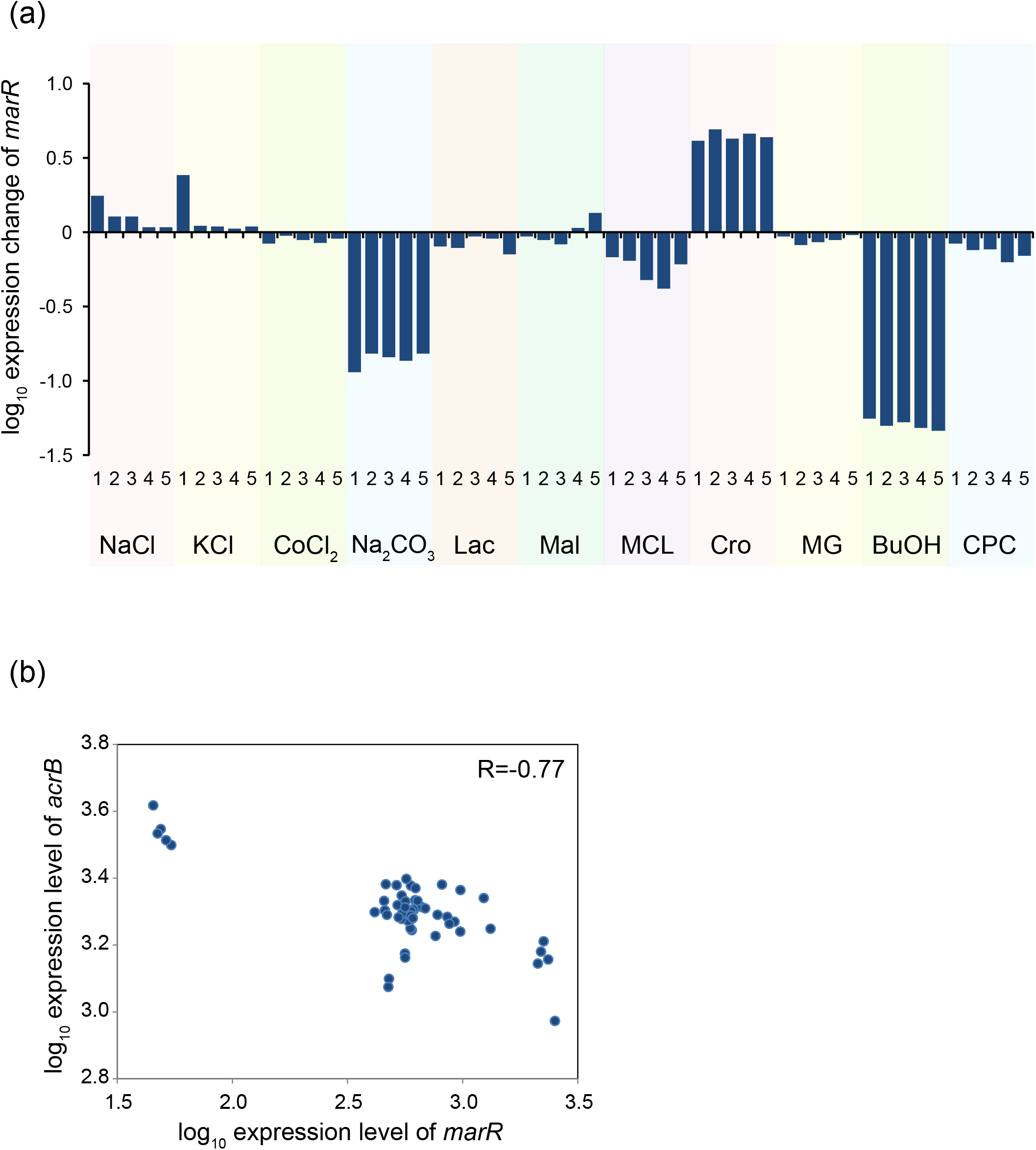
(a) Expression changes of *marR* in the resistant strains. (b) Correlation between expression levels of *marR* and *acrB*. Each dot represents the expression levels in each resistant strain.

**S2 Figure.**
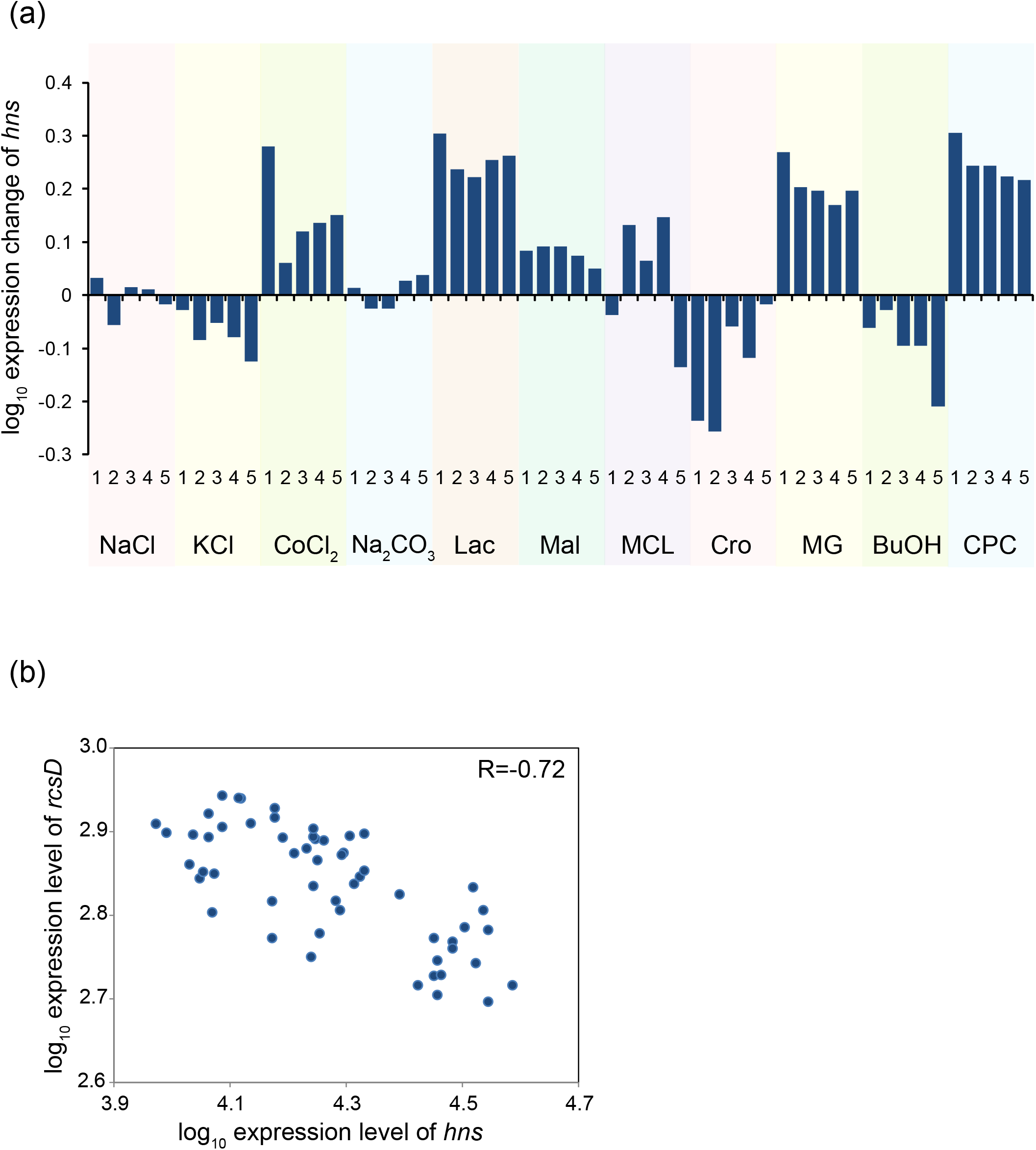
(a) Expression changes of *hns* in the resistant strains. (b) Correlation between expression levels of *hns* and *rcsD*.

**S3 Figure.**
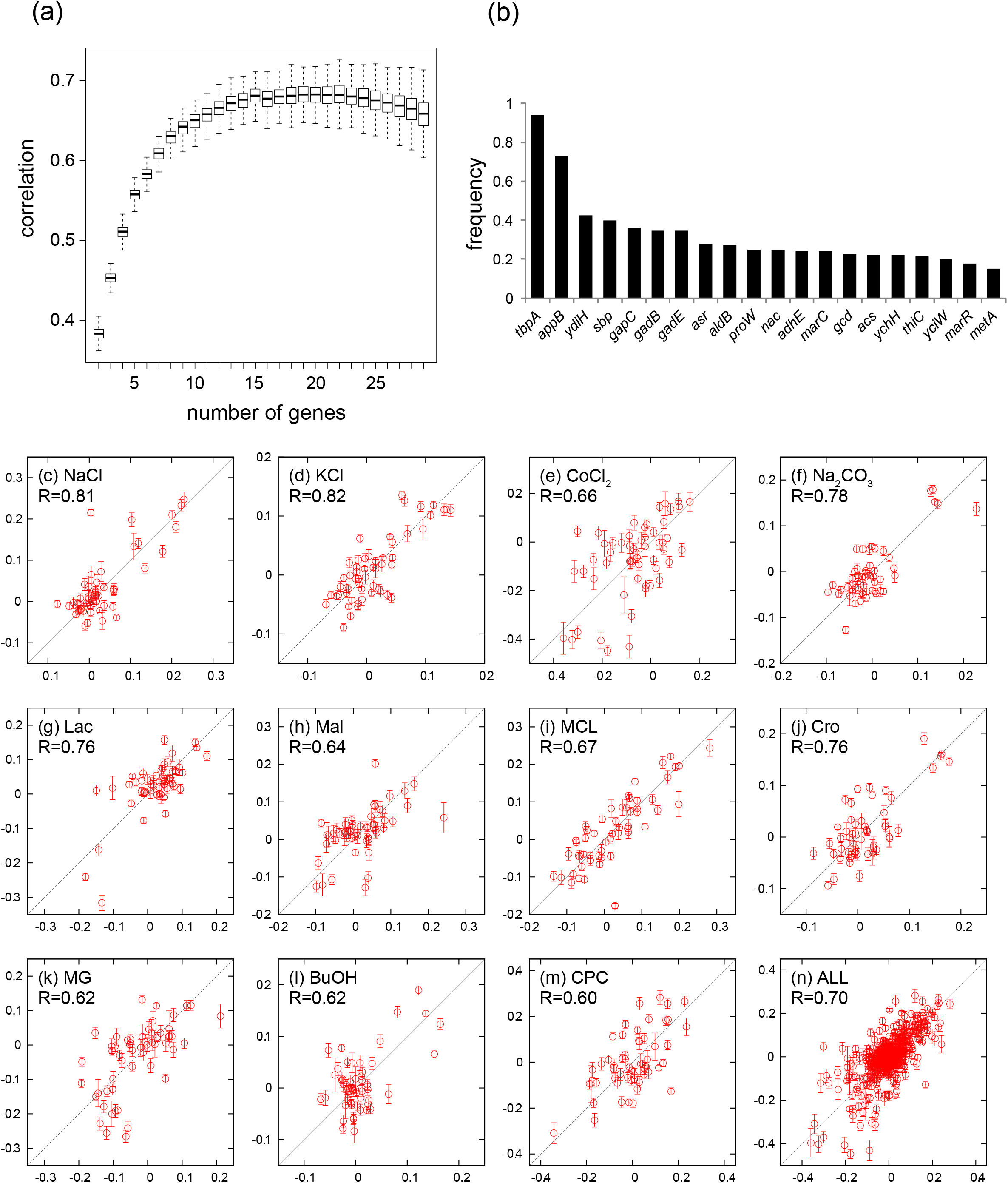
Prediction of stress resistance using transcriptomic data. (a) Prediction accuracy as a function of the number of genes, *N*, used for the fitting. The prediction accuracy was quantified by the correlation coefficients between predicted growth rate and observed growth rate of the test data. (b) Frequency of genes selected in GA trials in cases with *N*=15. Randomly generated 10,000 sets of test data and training data were used to calculate the frequency. Comparisons between observed and predicted growth rate for (c) NaCl, (d) KCl, (e) CoCl_2_, (f) Na_2_CO_3_, (g) Lac, (h) Mal, (i) MCL, (j) Cro, (k) MG, (l) BuOH, (m) CPC, and (n) all data were calculated by fitting using the following 15 genes: *tbpA, appB, ydiH, gadE, sbp, aldB, asr, marC, proW, tktB, nac, thiC, ydhZ, acs*, and *gcd*. Only test data not used for the fitting are plotted. The error bars in the y-axis were obtained by predicted growth rate calculated from 10,000 different sets of test data and training data.

**S4 Figure.**
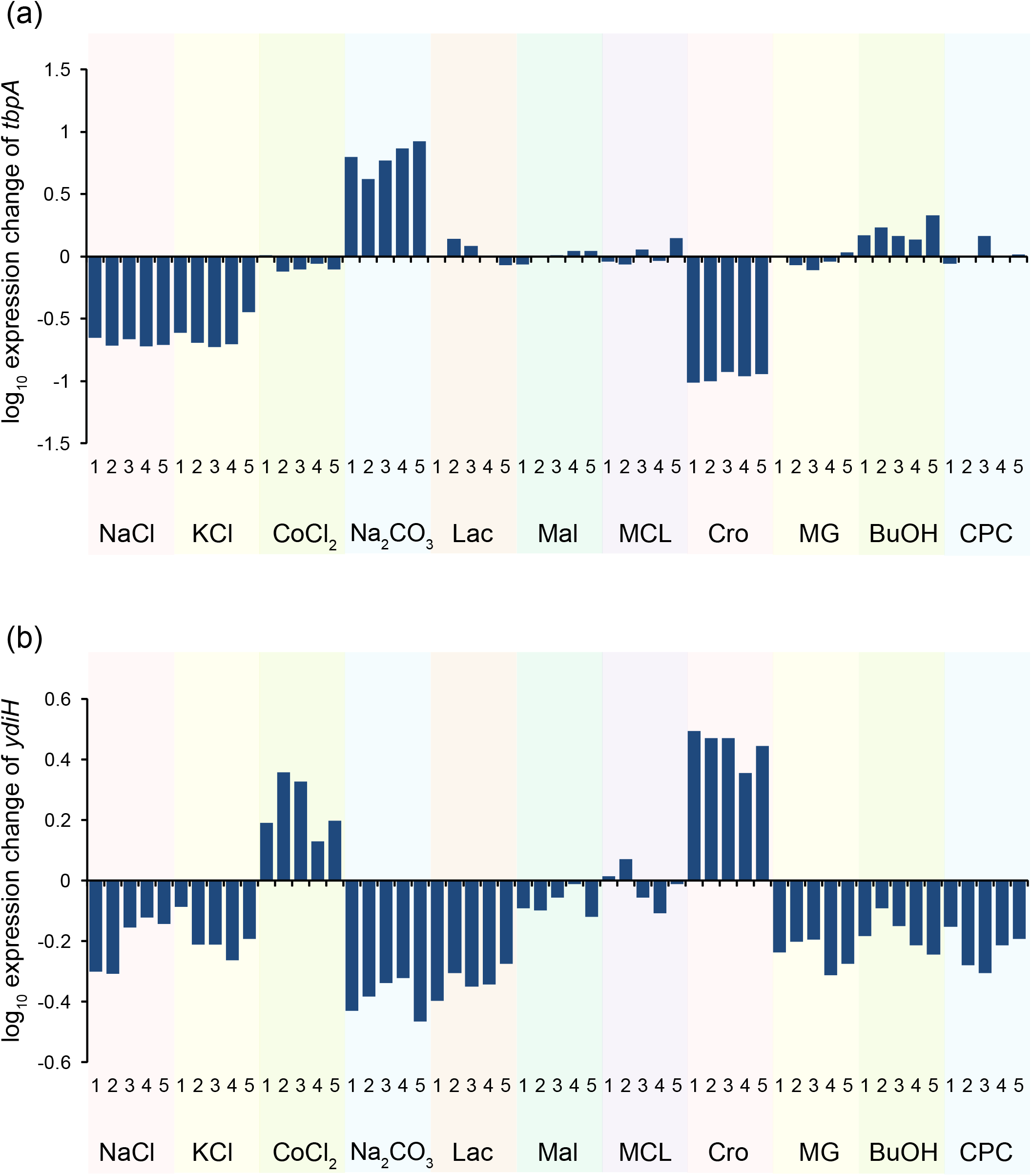
Expression changes of (a) *tbpA* and (b) *ydiH* in the resistant strains.

**S5 Figure.**
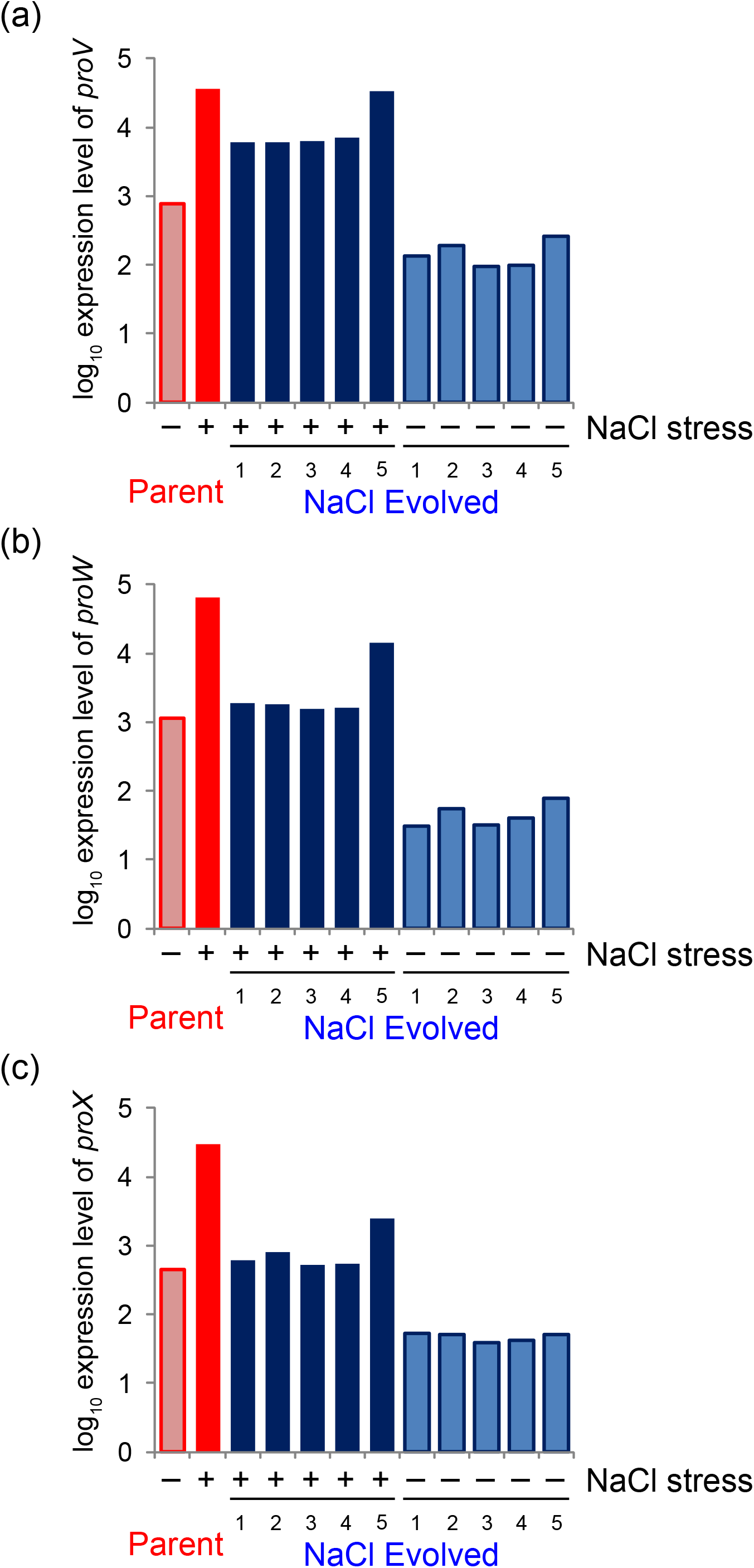
Expression levels of genes in *proVWX* operon. Log-transformed expression levels of (a) *proV*, (b)*proW*, and (c)*proX* in the parent and NaCl resistant strains with and without NaCl stress are plotted.

**S6 Figure.**
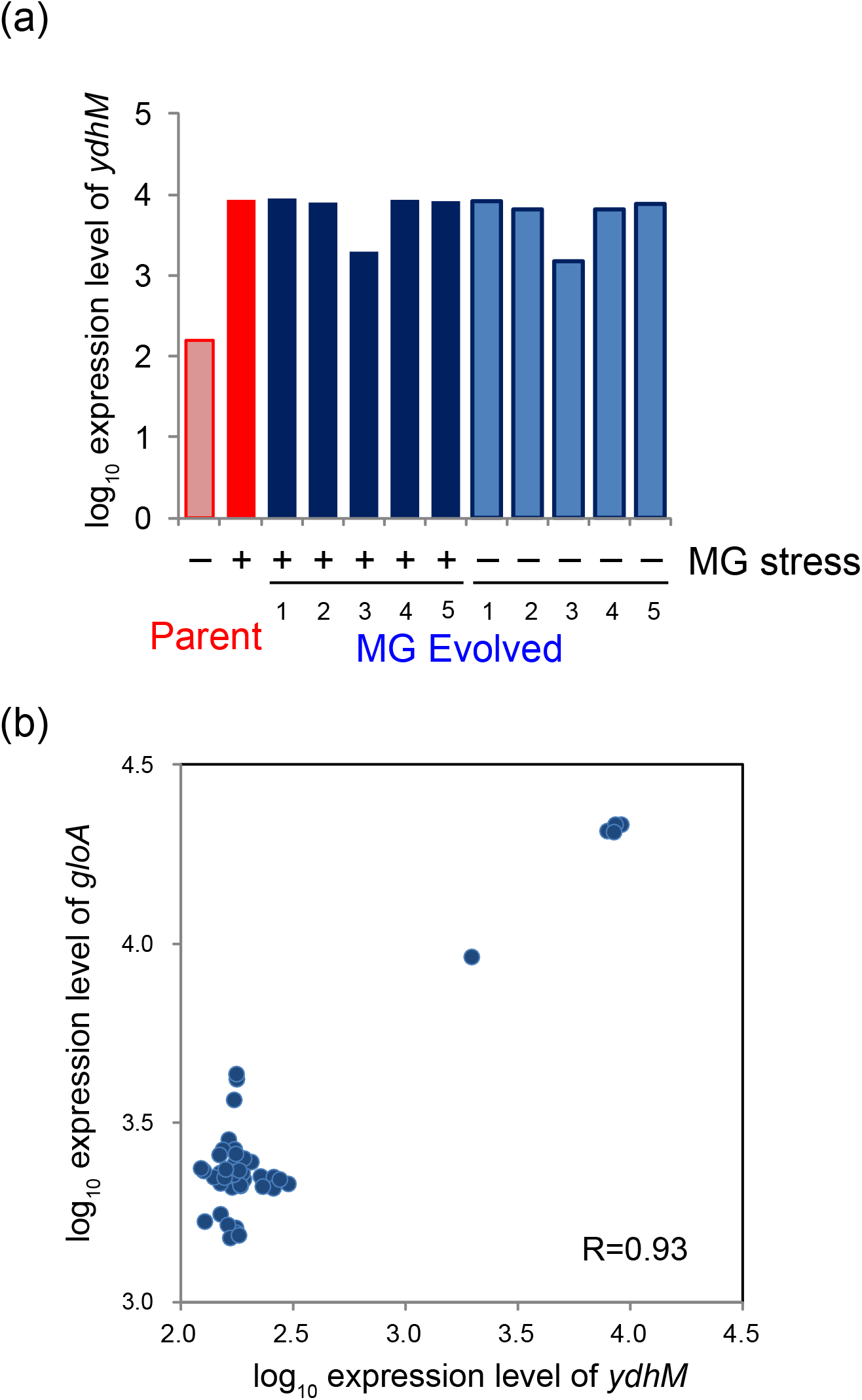
(a) Expression levels of *ydhM*. Log-transformed expression levels of *ydhM* in the parent and MG resistant strains with and without MG stress are plotted. (b) Correlation between expression levels of *ydhM* and *gloA*.

**S7 Figure.**
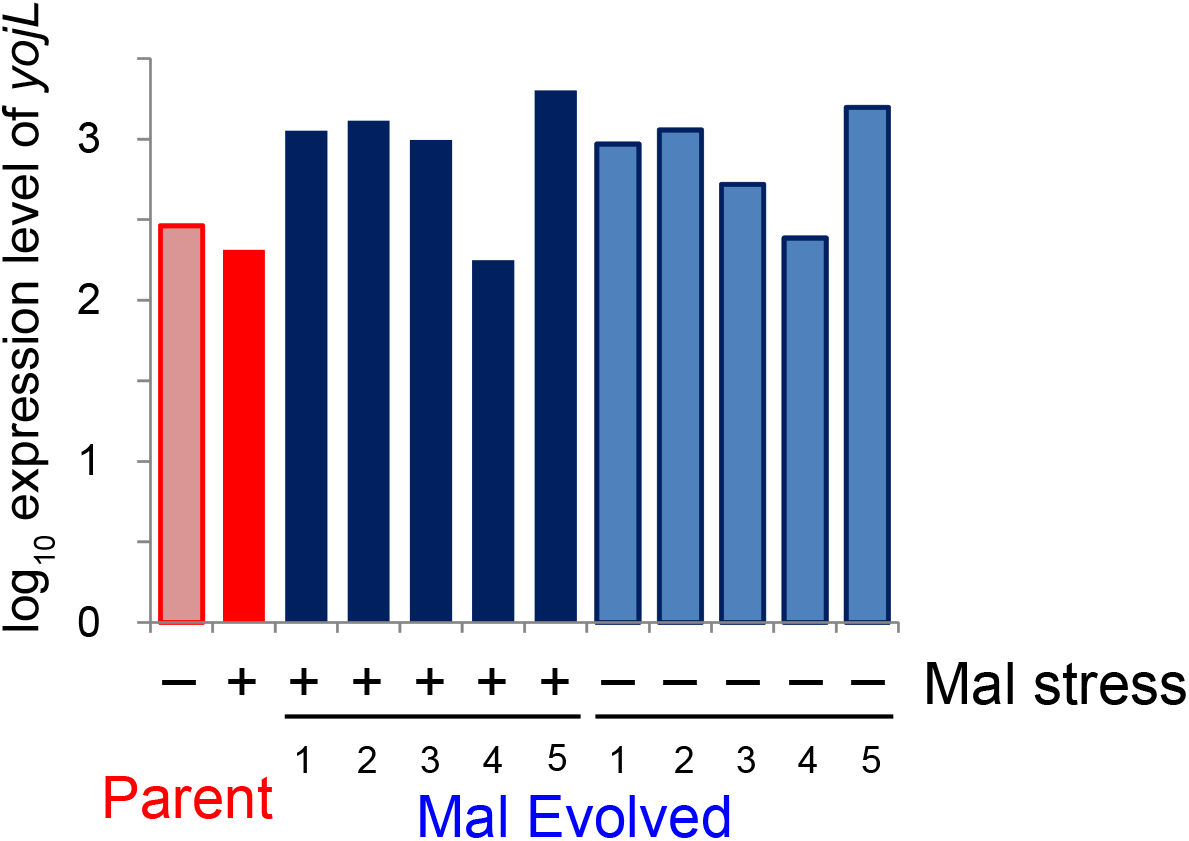
Expression levels of *yojL*. Log-transformed expression levels of *yojL* in the parent and Mal resistant strains with and without Mal stress are plotted.

**S8 Figure.**
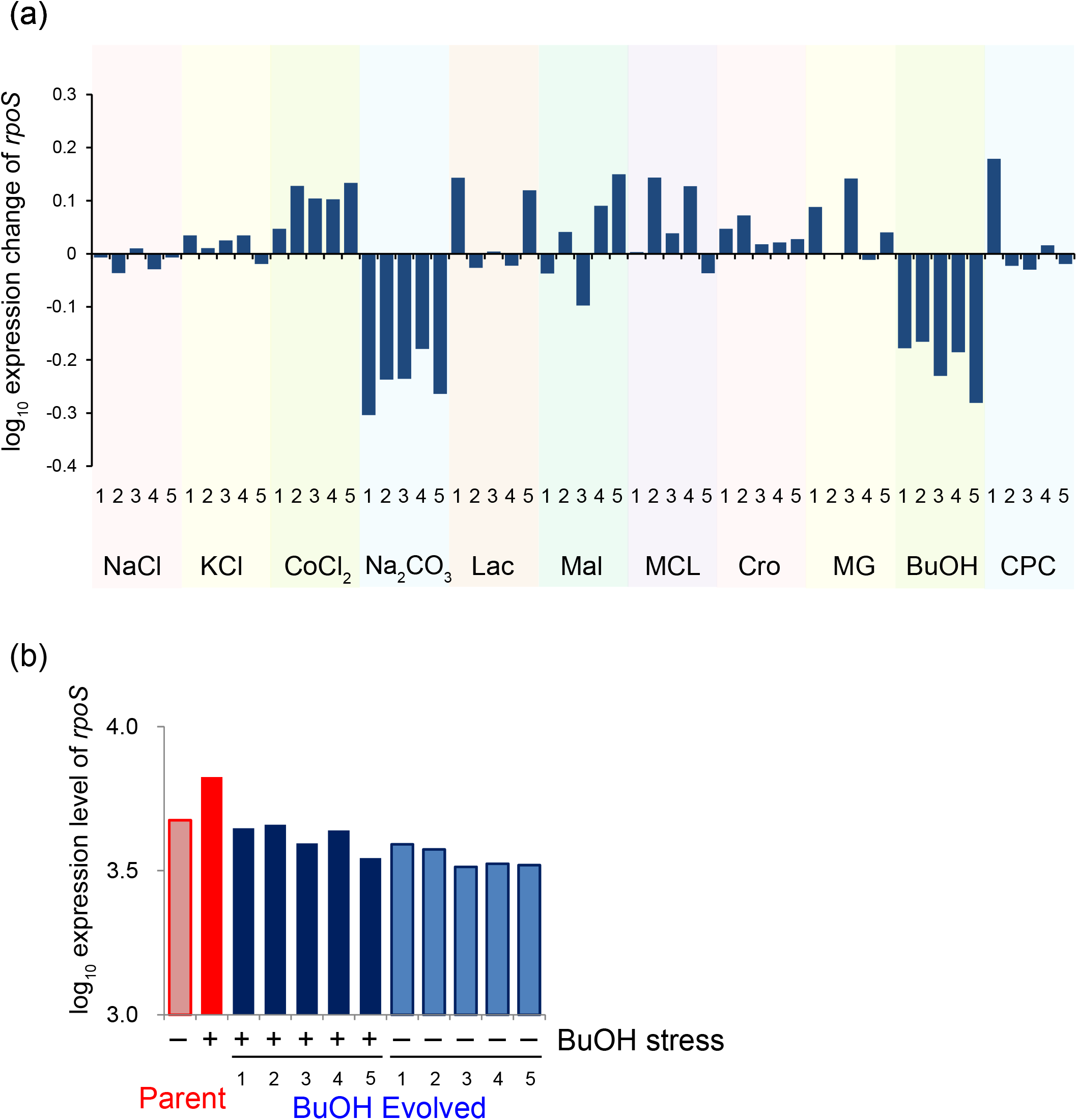
(a) Expression changes of *rpoS* in the resistant strains. (b) Expression levels of *rpoS* in the parent and BuOH resistant strains with and without BuOH stress are plotted.

**S9 Figure.**
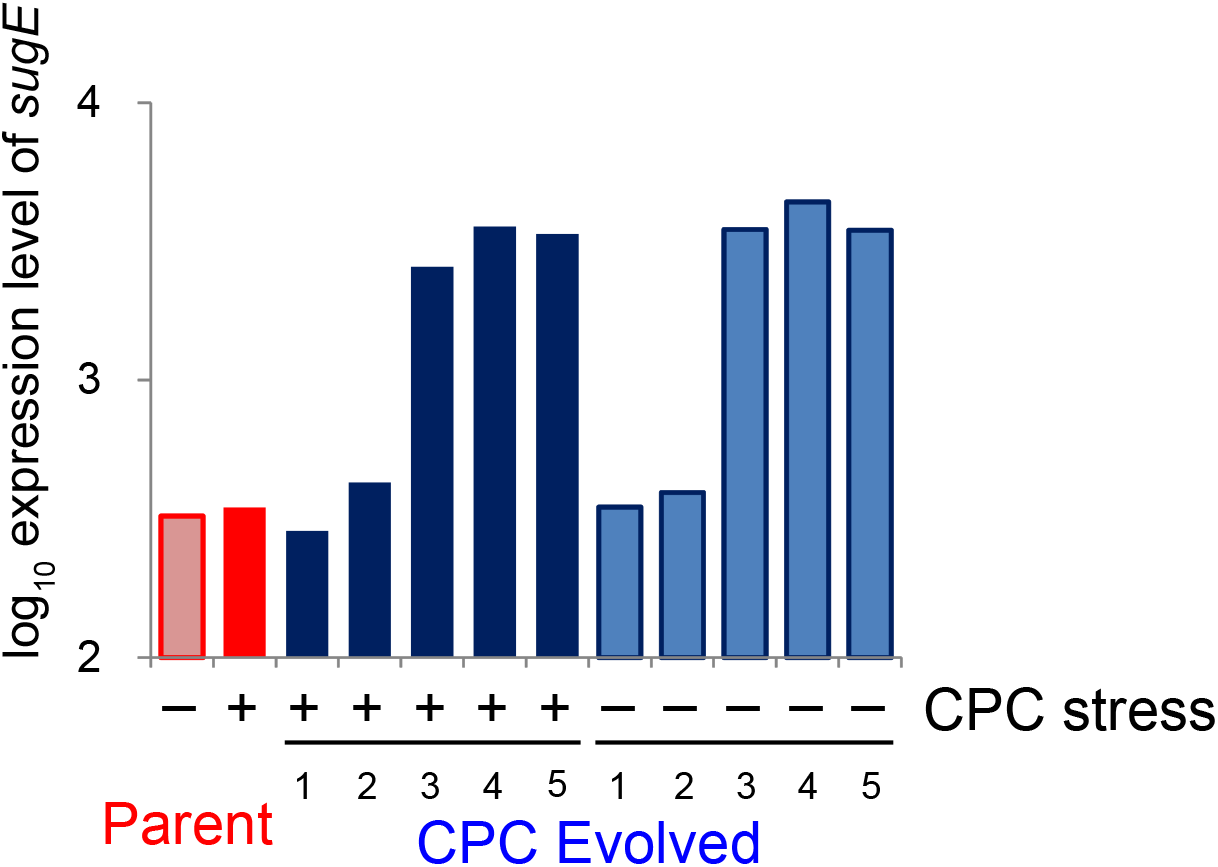
Expression levels of *sugE*. Log-transformed expression levels of *sugE* in the parent and CPC resistant strains with and without CPC stress are plotted.

**S1 Table.** Summary of relative growth rates of the resistant strains in stress conditions.

**S2 Table.** Transcriptome changes observed in the resistant strains.

**S3 Table.** List of mutations identified in the resistant strains.

## Supporting Text S1: Detailed description of common mutations identified in resistant strains

The number of mutations identified in the resistant strains is shown in Fig. 4, and detailed information about the mutations is presented in Supplementary Table S2. Below, for each stress we discuss the relationship between resistance acquisition, genome mutations and gene expression changes.

## Sodium chloride (NaCl) resistant strains

See Main Text.

## Potassium chloride (KCl) resistant strains

In KCl resistant strains, in addition to mutations in *proU* operon as described in Main Text, we found three resistant strains have mutations in the coding region of *ftsI*, which is involved in the cell division process. Since *ftsI* is an essential gene, we did not evaluate the effect of the identified *ftsI* mutations by introducing them into the parent strain. The role of *ftsI* mutations in KCl resistance remains unclear.

## Cobalt chloride (CoCl_2_) resistant strains

All resistant strains to CoCl_2_ stress had mutations in *corA*^1^, which encodes a transporter mediating influx of Mg^2+^, Ni^2+^, and Co^2+^. The identified mutations included frame-shift and deletion of ORF, suggesting that the disruption of *corA* activity contributed to CoCl_2_ resistance. This hypothesis was supported by the fact that the introduction of the *corA* mutation found in a CoCl_2_ resistant strain into the parent strain significantly increased the growth under CoCl_2_ stress (Fig. 5). Four out of five CoCl_2_ resistant strains also had mutations in *feoA/feoB* encoding ferrous iron transporters^2^. Even though the role of FeoA/FeoB in cobalt uptake has not been demonstrated^3^, these identified *feoA/feoB* mutations suggested that FeoA/FeoB transporters are involved in Co^2+^ transport.

## Sodium carbonate (Na_2_CO_3_) resistant strains

Two out of five Na_2_CO_3_ resistant strains had mutations in *sapABCDF* operon, which encode Sap (Sensitive to antimicrobial peptides) ABC importer^4^. To evaluate the effect of the mutations, we introduced *sapA* mutation in its coding region (found in Na_2_CO_3_-2 resistant strain) into the parent strain, and confirmed that it significantly increased the growth under Na_2_CO_3_ stress. Sap transporter is known to contribute alkali stress resistance in *Sinorhizobium meliloti*, root nodule bacteria^5^, and thus our result might suggest that Sap transporter is involved in alkali stress resistance also in *E. coli*.

## L-Lactate (Lac) resistant strains

Four Lac resistant strains (Lac-1, 2, 3, and 4) had mutations either *purT* or *purU* genes, which are involved in purine biosynthesis^6^. In *Lactobacillus lactis*, it was suggested that perturbing purine biosynthesis is related to multi-stress resistance by changing 3’,5’-bispyrophosphate (ppGpp) concentration^7^, which is known as a master regulator of the stringent response to amino acid starvation. Thus, the identified mutations in *pur* genes might be related to the control of the stringent response, although the details still remain unclear. We confirmed that the introduction of the mutation in the *purT* coding region significantly increased the growth under Lac stress (Fig. 5).

## L-Malate (Mal) resistant strains

Four Mal resistant strains (Mal-1, 2, 3, and 5) had mutations in the promoter region of *yojL* (*apbE*). The expression levels of *yojL* in these four strains significantly increased (Fig. S7), probably due to the mutations in the promoter region. *yojL* encodes a periplasmic lipoprotein which is involved in thiamine biosynthesis^8^, and also it is thought to play a role in assembly or maintenance of iron-sulfur clusters^9^. These mutations commonly fixed in *yojL* promoter region suggests their contribution to Malate resistance by up-regulating its expression level. However, we could not observe a significant growth rate increase under Mal stress when one of these mutations was introduced in the parent strain (Fig. 5).

## Methacrylate (MCL) resistant strains

All resistant strains to MCL had mutations in *pykF*, which encodes a pyruvate kinase that catalyzes the conversion of phosphoenolpyruvate (PEP) into pyruvate in the central metabolic pathway. All these mutations in *pykF* were nonsynonymous single nucleotide substitutions, which might suggest that the change of enzymatic activity contribute to the methacrylate resistance. To verify this, we introduced the mutation found in a MCL-3 resistant strain into the parent strain, and confirmed that it significantly increased the growth rate under methacrylate stress (Fig. 5). Although the mechanism how *pykF* mutations contribute to MCL resistance remains unclear, controlling intracellular accumulation of pyruvate caused by MCL stress might be related to the mechanism of resistance. MCL is known to inhibit the enzymatic activity of pyruvate formate-lyase (Pfl) that converts pyruvate into acetyl-CoA and formate^10^, and thus MCL stress can cause an increase in intra-cellular pyruvate. Since the PEP/pyruvate concentration ratio is an important parameter to control various metabolic pathways, including glucose uptake via sugar phosphotransferase system (PTS)^11^, the decrease of PykF activity by the mutations might play a role in rebalancing the disrupted PEP/pyruvate ratio by MCL stress.

## Crotonate (Cro) resistant strains

No common mutation was identified in Cro resistant strains.

## Methylglyoxal (MG) resistant strains

See Main Text.

## *n*-butanol (BuOH) resistant strains

Three BuOH resistant strains (BuOH-3, 4, and 5) had mutations in the coding region of *cspC*, and we confirmed that one of these mutations can slightly increase the BuOH stress resistance (Fig. 5). CspC is a constitutively produced member of the CspA family of RNA-binding proteins, which increases RpoS expression level presumably by stabilizing its mRNA^12^. RpoS is a central regulator of the general stress response whose up-regulation leads to growth arrest^12^. We found that the *rpoS* mRNA expression increased in response to BuOH stress addition, while it relaxed to the original levels in BuOH resistant strains (Fig. S8). This result suggests that the growth rate increase under BuOH stress can be partially explained by the down-regulation of *rpoS* caused by the *cspC* mutations. Furthermore, all BuOH resistant strains had deletion of region including *yneK, ydeA*, and *marC* genes. Interestingly, disruption of *marC* was also identified in isobutanol resistant *E. coli* strains obtained by laboratory evolution^13,14^. Although *marC* is a poorly characterized gene whose function is unknown, these results suggest a positive effect of *marC* disruption on butanol resistance.

## Cetylpyridinium chloride (CPC) resistant strains

Three out of five CPC resistant strains (CPC-2, 4, and 5) had mutations in *dcm*, which encodes a DNA cytosine methyltransferase. To verify the effect of *dcm* mutations, we introduced the nonsynonymous single nucleotide substitutions found in CPC-5 strain into the parent strain. This mutant strain exhibited a slight increase of growth rate under the CPC stress (Fig. 5), although the difference was not statistically significant. This result might suggest that the mutation in *dcm* has a weak fitness gain effect in the CPC stress environments. The mechanism for CPC resistance by these *dcm* mutations remains unclear. A recent study demonstrated that the deletion of *dcm* contributes to antibiotic resistance through up-regulation of *sugE* encoding multidrug efflux transporters^15^. In three of five CPC resistant strains (CPC-3, 4, and 5), mutations are found in the promoter region of *sugE*, which suggests that these resistant strains acquired CPC resistance by activating the SugE efflux transporter. The fact that the expression levels of *sugE* significantly increased in CPC-3, 4, and 5 strains and weakly increased in CPC-2 strain supported this hypothesis (Fig. S9).

